# SCSA: a cell type annotation tool for single-cell RNA-seq data

**DOI:** 10.1101/2019.12.22.886481

**Authors:** Yinghao Cao, Xiaoyue Wang, Gongxin Peng

**Affiliations:** Center for Bioinformatics, Institute of Basic Medical Sciences, Chinese Academy of Medical Sciences, School of Basic Medicine, Peking Union Medical College, Beijing, China

**Author notes:** Corresponding Author Xiaoyue Wang, Gongxin Peng.

**Keywords:** single-cell RNA sequencing, cell type annotation, CellMarker database, score annotation model, differentially expressed genes

## Abstract

Currently most methods take manual strategies to annotate cell types after clustering the single-cell RNA sequencing (scRNA-seq) data. Such methods are labor-intensive and heavily rely on user expertise, which may lead to inconsistent results. We present SCSA, an automatic tool to annotate cell types from scRNA-seq data, based on a score annotation model combining differentially expressed genes (DEGs) and confidence levels of cell markers from both known and user-defined information. Evaluation on real scRNA-seq datasets from different sources with other methods shows that SCSA is able to assign the cells into the correct types at a fully automated mode with a desirable precision.

## Introduction

Recent development of scRNA-seq methods has enabled unbiased, high-resolution transcriptomic analysis of individual cells in a heterogeneous cell population (Tang et al., 2009; Picelli et al., 2013a; Kolodziejczyk et al., 2015; Haque et al., 2017). scRNA-seq methods have been used to characterize thousands to millions of cells in developing embryos (Chu et al., 2016), immune cells (Shalek et al., 2013), complex tissues such as brain (Zhong et al., 2018) and tumor (Chung et al., 2017), which have greatly promoted our understanding of human development and diseases.

At the core of myriad scRNA-seq applications is the ability to identify different cell types and cellular states from a complex cell mixture based on gene expression profiles. Recently, several computational methods have been developed for annotating cell types in scRNA-seq using cell-based mapping strategy. For instance, SingleR (Aran et al., 2019) infers the cell type for each of the single cells using a novel hierarchical clustering method based on similarity to the reference transcriptomic datasets of purified cell types. Similarly, scMatch (Hou et al., 2019) annotates single cells by identifying their closest match in gene expression profiles of large reference dataset, such as the Functional Annotation Of The Mammalian Genome 5 (FANTOM5) resource (Brown et al., 2009; Lizio et al., 2017). However, such approaches required transcriptome of purified cells or pre-annotated scRNA-seq data, ideally under the same experimental design using the same platform, which is often not available. Using prior knowledge of cell-type specific marker genes increased the accuracy and efficiency of cell type assignment, allowing for identification of both known and *de novo* cell types in scRNA-seq data of complex tissues, as shown by CellAssign (Zhang et al., 2019a) and Garnett (Pliner et al., 2019a).

A more common practice for cell type annotation is the cluster-then-annotate approach. Unsupervised clustering methods based on dimension reduction algorithms such as principal component analysis (PCA) and t-distributed stochastic neighbor embedding (t-SNE) have been developed to partition the cells based on the similarity of their gene expression patterns (Bacher and Kendziorski, 2016; Kiselev et al., 2019). And users could manually assign a cell type to each cluster based on differentially expressed markers by consulting the literature for cell-type specific gene markers. However, there is still some problem associated with the manual annotation step. For example, the canonical marker genes used in the assignment process may have an impact on the annotation accuracy, which may lead to biased results with uncontrolled vocabularies for cell type labels in different datasets. Expert-curated knowledge databases such as CellMarker (Zhang et al., 2019b) and CancerSEA (Yuan et al., 2019b), have been developed to provide a comprehensive and unified resource of cell markers for various cell types in human and mouse tissues. Yet it is still lack a method to leverage the information in these databases for annotation. Furthermore, marker genes could express in more than one cell type, make the annotation more complex for human to process.

To overcome these difficulties and to streamline the cell type assignment process for scRNA-seq data, we developed SCSA, an algorithm that can automatically assign cell types for each cell cluster in scRNA-seq data. SCSA uses marker genes of known cell types highly expressed in a cell cluster to label the cluster. It can be directly applied to clustering results generated from other scRNA-seq analysis softwares such as CellRanger (Zheng et al., 2017) and Seurat (Butler et al., 2018). To eliminate the bias of marker selection, SCSA integrates all marker genes to cell-type matrix from CellMarker (Zhang et al., 2019b) and CancerSEA (Yuan et al., 2019a) database, using a score annotation model that accounts for quantitative information and discrepancies information of marker genes. For cell clusters lacking known cell markers, SCSA will also perform a Gene Ontology (GO) enrichment analysis and report the results to give some clues to the user. Through extensive evaluation on several real scRNA-seq datasets of both human and mouse origin generated from different platforms including Smart-seq (Ramskold et al., 2012),Smart-seq2 (Picelli et al., 2013b) and 10x Genomics (Zheng et al., 2017). We demonstrated that SCSA has an excellent and unbiased performance on cell type annotation with a desirable precision.

## Materials and methods

### Marker genes identification

The input of SCSA is a DEGs clusters matrix, in a format that is supported by the clustering output of CellRanger or Seurat. Based on the matrix, SCSA identifies the marker genes of each cell cluster through a filtration with log2-based fold-change (LFC) value and P-value (LFC >= 1, P <= 0.05). For each cell cluster, a marker gene identification vector is generated for j genes with LFC values, which is defined as *E*_*j*_ *=* {*e*_1_,*e*_2_,⋯*e*_*j*_} *=* (*e*) _*j*×1_, here, *e* represents the absolute value of LFC multiplied by mean of all.

### Cell marker database

In order to improve the accuracy of cell cluster annotation for scRNA-seq data, SCSA uses cell markers from two public databases: CellMarker and CancerSEA. SCSA integrated 11,464 manually curated cell markers of 459 cell types in 158 human tissues/sub-tissues and 7,855 cell markers of 385 cell types in 80 mouse tissues/sub-tissues from CellMarker database. Also SCSA integrates 1,244 markers from CancerSEA database, which consists of 14 functional states from 25 human cancer types. Furthermore, SCSA can accept user-defined marker gene database as additional information for cell cluster annotation. The user-defined marker gene database must have two columns, with the name of cell types in the first column and marker gene for each cell type in the second column. In that case, SCSA will combine both known databases and the custom database to predict the annotations for cell clusters.

### Annotation model construction

For those genes which existed in both the DEGs and known cell marker databases, SCSA constructs a cell-gene sparse matrix (defined as 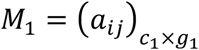 with *c*_1_(*c*_1_ ≤ *i*) cells and *g*_1_(*g*_1_ ≤ *j*) genes as “marker evidence”. Here, for each cell *i* and each gene j in the matrix *M*_1_, *a* refers to the sum number of references in the CellMarker database. To eliminate the huge differences of marker evidence between the well-known gene and less-known genes, we transform the value to log2-based and plus a constant (0.05). Also, to represent the whole gene set for a certain cell, we create a cell type style vector which takes multiplication of standard deviation of the marker evidence and marker numbers (defined as *L*_1_ *=* {*l*_1_,*l*_2_,⋯*l*_*c*1_}, where *l = std*(*a*_*ij*_) * *num*(*a*_*ij*_>0)). So, for the known marker database, the raw score vector of a cell type is 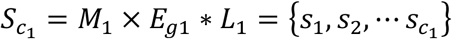.

For marker databases from multiple sources including known and user-defined ones, with *k* as the total number of databases, multiple cell-gene sparse matrices could be constructed according to the DEGs, which are defined as *M*_*k*_ *=* (*a*_*ij*_)_*p*×*q*_, *p* × *q* ∈{(*c*_1_ × *g*_1_),(*c*_2_ × *g*_2_),…,(*c*_*k*_ ×*g*_*k*_)}. With the definition of multiple gene expression vectors defined as *E*_*k*_ *=* (*e*)_*q*×1_,*q* ∈{*g*_1_,*g*_2_,⋯,*g*_*k*_},*q* ≤ *j*), *k* raw score vectors will be generated, which is defined as *S*_*k*_ *= M*_*k*_ *E*_*k*_ *L*_*k*_ *=* (*a*_*ij*_) _*p*×*q*_ × (*e*)_*q*×1_ * (*l*)_*q*×1_ *=* (*s*)_*p*×1_,*p* ∈{*c*_1_, *c*_2_,⋯*c*_*k*_}. Suppose *S*_*k*_ to be a function to the score vectors, which is defined as *F*_*l*_. To eliminate the difference of those vectors and compare them with each other standardly, SCSA performs z-score normalization for them. In detail,

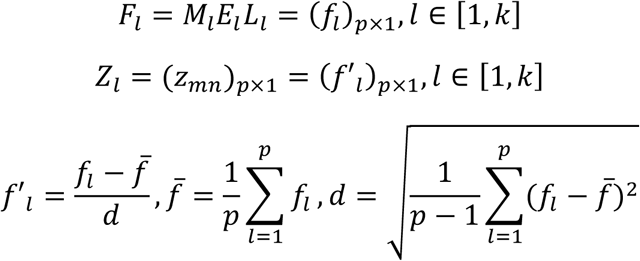

Notably, the score vectors derived from different kinds of databases may have different lengths. To give a uniform score to a certain cell type, SCSA transforms them to the same length:

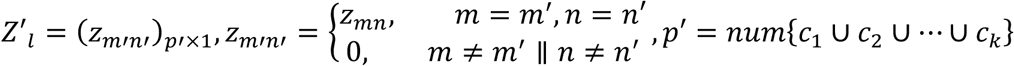

Finally, an annotation model will be constructed by merging the database weight coefficient matrix *W* and the last uniform score vector.

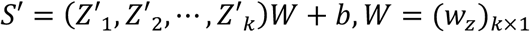

### GO enrichment analysis

Not all gene markers of cell types have been curated in known cell marker databases. To solve this problem, SCSA employs a GO enrichment analysis to give some clues to the user to identify new cell types. In detail, for a certain GO term, SCSA uses Fisher’s exact test to calculate the P-values, using DEGs of the selected cluster as foreground values and DEGs in other clusters as background values, respectively. After that, P-value was adjusted by the Benjamini-Hochberg (BH) method (Benjamini and Hochberg, 1995).

### FANTOM5+SingleR dataset

FANTOM5+SingleR dataset was collected from GitHub repository of scMatch (https://github.com/forrest-lab/scMatch). The dataset contained gene expression data of 916 human samples (660 primary cell samples and 256 cell lines) and 821 mouse samples (471 primary cell line samples, 302 tissue samples, and 48 transformed cell lines) from FANTOM5 and 972 human samples and 1188 mouse samples in SingleR’s reference dataset. Annotation of these cells into certain categories were also downloaded from the same repository. For human cells in FANTOM5+SingleR dataset, terms of cell types associated with more than 10 samples were collected as the FANTOM5_hs1 dataset. And cell types associated with more than 40 samples were used FANTOM5_hs2 datasets. Similarly, FANTOM5_mm1 contained cell types with more than 10 samples and FANTOM5_mm2 included cell types with more than 40 samples in the FANTOM5+SingleR mouse dataset.

### Real scRNA-seq datasets

Four real datasets (GSE72056 (Tirosh et al., 2016b), GSE81861 (Li et al., 2018), E-MTAB-6149 (Lambrechts et al., 2018), and the PBMCs dataset were used for evaluation of SCSA. GSE72056, GSE81861 and E-MTAB-6149 datasets are single cell data obtained from human tumors (melanoma, colorectal and lung) using Smart-seq2, Smart-seq and 10X genomics technology, respectively. Normal cells in the GSE72056 dataset was downloaded from Gene Expression Omnibus (GEO) repository (https://www.ncbi.nlm.nih.gov/geo/query/acc.cgi?acc=GSE72056) and used to optimize the parameters for SCSA. Normal cells in the GSE81861 dataset was downloaded from GEO (https://www.ncbi.nlm.nih.gov/geo/query/acc.cgi?acc=GSE81861). E-MTAB-6149 was downloaded from ArrayExpress repository (https://www.ebi.ac.uk/arrayexpress/experiments/E-MTAB-6149).

The peripheral blood mononuclear cells (PBMCs) datasets, including four scRNA-seq data of peripheral blood mononuclear cells (3k, 4k, 6k and 8k of PBMCs) were downloaded from the 10X Genomics official website (https://www.10xgenomics.com/resources/datasets). The 3k and 6k data were blood samples from one donor generated using the v1. chemistry and preprocessed with CellRanger1.1.0. They were labeled “3k PBMCs from a healthy donor” and “6k PBMCs from a healthy donor”, respectively. The 4k and 8k datasets, under the label “4k PBMCs from a healthy donor” and “8k PBMCs from a healthy donor”, were samples collected from one donor, generated using the v2. Chemistry and preprocessed using CellRanger2.1.0.

### Evaluation of SCSA performance on real datasets

For the known cell type datasets, we defined cell type cluster by their real cell types. Then we generated calculated the DEGs for each clusters using two methods. One is traditional Student’s t-test perform by in-house script and the other using Seurat. We obtained the final two lists of DEGs for the input of SCSA through setting the threshold with P-value of 0.001 and LFC value of 1.

For the 4 PBMCs datasets, data preprocessing, normalization and unsupervised clustering were already performed by CellRanger workflow from its website. Since 10X Genomics official website illustrated a workflow example using 3k PBMCs from a healthy donor containing 5 cell clusters and gave a final annotation results contained monocytes, T cells, NK cells, megakaryocytes, and B cells, respectively, we choose results with five clusters to do the further evaluation. SCSA identified the DEGs of each cluster through the LFC (LFC>=1.5) value and P-value (P<=0.05) and predicted the cell types according to the clusters.

To evaluate the stability of SCSA in annotating the cell type of a cluster, a heat map was generated using hierarchical clustering method for all cell types of top five scores in a cell cluster. To further demonstrate the accuracy of SCSA, we calculated the percentage of the five clusters cell types to measure the abundance of the same cell type based on the prediction of SCSA in the four PBMCs datasets, respectively.

### Comparison with other cell type annotation tools

Three annotation tools, scMatch (Hou et al., 2019), CellAssign (Zhang et al., 2019a) and Garnett (Pliner et al., 2019a) were used for comparison. scMatch was run with default parameters using FANTOM5 as the reference dataset. For CellAssign (version 0.99.2), a build-in set of markers contain 8 common cell types in human tumor microenvironment were used to test for human datasets. For Garnett (version 0.1.14), two pre-trained classifiers, trained from human PBMCs tissue (Garnett_human_pbmc) and lung tissue (Garnett_human_lung), were downloaded from (https://cole-trapnell-lab.github.io/garnett/classifiers) and used for classification. Because CellAssign and Garnett only annotate cell types for each of single cells instead of cell cluster, to make the evaluation at the same level, all cells of a certain cluster were predicted and the cell type with the maximum occurrence was chosen as the final predicted result. Three tags were used to compare the results. “Positive”meant correct prediction to the cell type. “Negative” meant incorrect prediction cell type. “Missed” meant no clear cell type prediction. Accuracy of cluster was calculated as sum of clusters with “Positive” tag divided by sum of all clusters.

### Software availability

SCSA is implemented in Python3 as an open source software under the GNU General Public License, and the source code is freely available together with full documentation at https://github.com/bioinfo-ibms-pumc/SCSA.

## Results

### Design principles of SCSA

The SCSA algorithm is a three-step procedure that includes marker genes identification, annotation model construction, and GO enrichment analysis (Figure 1). First, the input of SCSA is a gene expression matrix with cell cluster information (such as the output results of CellRanger or Seurat). SCSA identifies a group of marker genes for each cluster from input expression matrix by differential gene expression analysis. Next, for each cluster, genes identified as marker genes that have one or more linked cell types in a database will be used to generate a cell-gene matrix for that cluster. For each cell type in the matrix, SCSA then used a decision model to assign a score by combining the enrichment of marker gene expression and the strength of evidence for the marker genes in the database. SCSA could also take marker gene information from multiple databases and assign different weights to them. Finally, SCSA provides an alternative gene ontology enrichment analysis step to give some clues to the user for the function of a cell cluster in addition to its annotation.

**Figure 1.**
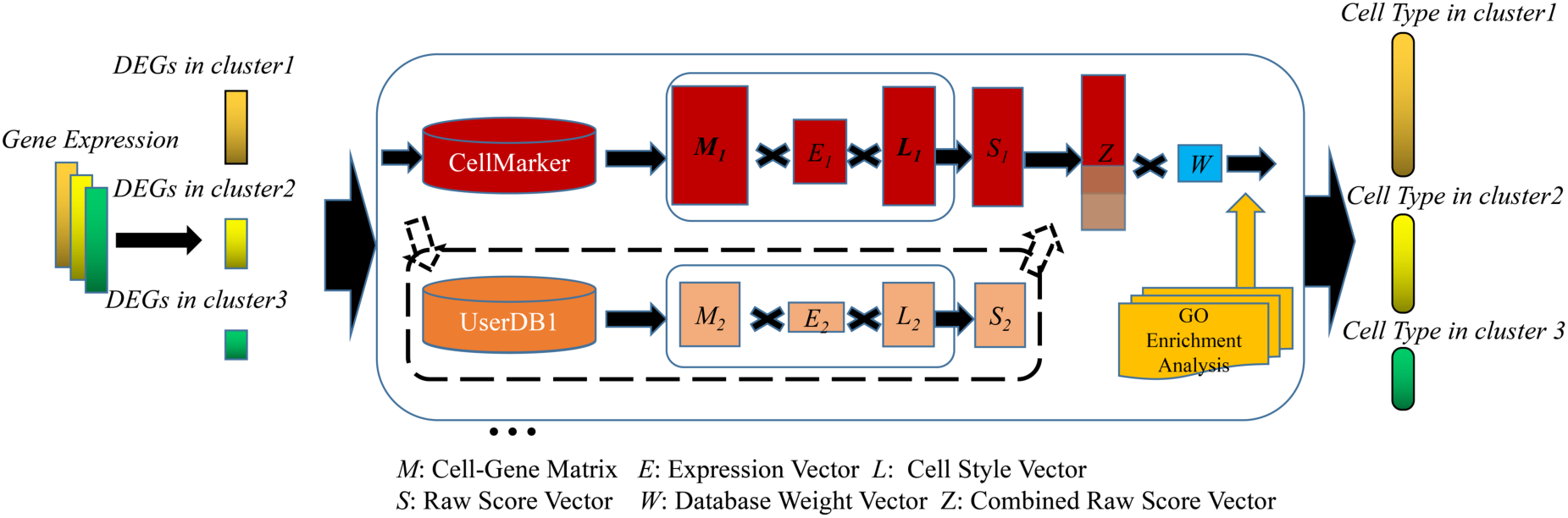
Flowchart of the SCSA. First, DEGs of each cluster will be extracted and filtered from gene expression file. Next, SCSA employs marker gene databases to annotate cell clusters. In this step, both known marker gene database and user-defined marker database could be used simultaneously. For each cluster each database, a cell-gene matrix (*M*) with two vectors (*E, L*) will be generated to form a raw score vector (*S*). If multiple databases were selected, vectors would be normalized and combined together to make a new vector (*Z*), then multiplied with a database weight matrix (*W*) to make the last uniform vector. In the last step, ranked cell type vector will be generated according to the uniform score. In addition, SCSA employs GO enrichment analysis to give users some clue for unidentified clusters.

### Evaluation performance of SCSA on known datasets

We first evaluated the performance of SCSA on three scRNA-seq datasets with known cell types (Table S1). The GSE72056 dataset include 2,840 single cells isolated from normal tissues of 19 melanoma patients (Tirosh et al., 2016a). These cells have been annotated manually by experts into six cell types, namely B cells, T cells, Cancer-associated fibroblast (CAF), Macrophage, Endothelial cells and Natural killer cells. SCSA accurately annotated five of the six cell types, except for the “Cancer-associated fibroblast (CAF)” cells (Table S2, S3). Instead, SCSA gave the CAF cluster a different label named “Mesenchymal stem cell”. It has been reported that fibroblasts shared more common features with mesenchymal stem cells (Haniffa et al., 2009) by expressing similar cell immunophenotypic markers, as well as the genes that are known to be expressed in stem cells (Brohem et al., 2013). Therefore, we considered the “Mesenchymal stem cell (MSC)” label SCSA reported concordant with the “CAF” label assigned by human expert. Similarly, for seven known clusters in GSE81861 dataset (Li et al., 2017), which contains 265 single cells derived from normal tissues adjacent to 11 colorectal tumors, SCSA correctly predicted six of the seven clusters and identified the “Fibroblast” group as “MSC” (Table S2, S3). In the lung cancer dataset of 45,232 normal cells containing 7 known cell types (Lambrechts et al., 2018), SCSA accurately assigned B cells, Endothelial cells, Epithelial cells, Myeloid and T cells, and identified “Fibroblast” group as “MSC” and “Alveolar” as “Epithelial Cell” (Table S2, S3).

In order to evaluate the robustness of SCSA on large scRNA-seq datasets, we used four PBMCs (3k, 4k, 6k and 8k) datasets from 10X genomics website. We collected all possible cell types of a cell cluster according to the top five scores under the score annotation model of SCSA. The correlation of all cell types and scores were calculated and compared. As shown in Figure 3A, based on the five annotated cell types (monocyte cells, T cells, NK cells, megakaryocytes cells, and B cells) of CellRanger, SCSA achieved a great consistency in the four PBMCs datasets. Notably, the “macrophage cell” type was predicted as the second top score for the cluster, which was annotated as “monocytes” by SCSA due to the reason that they share many marker genes.

**Figure 2.**
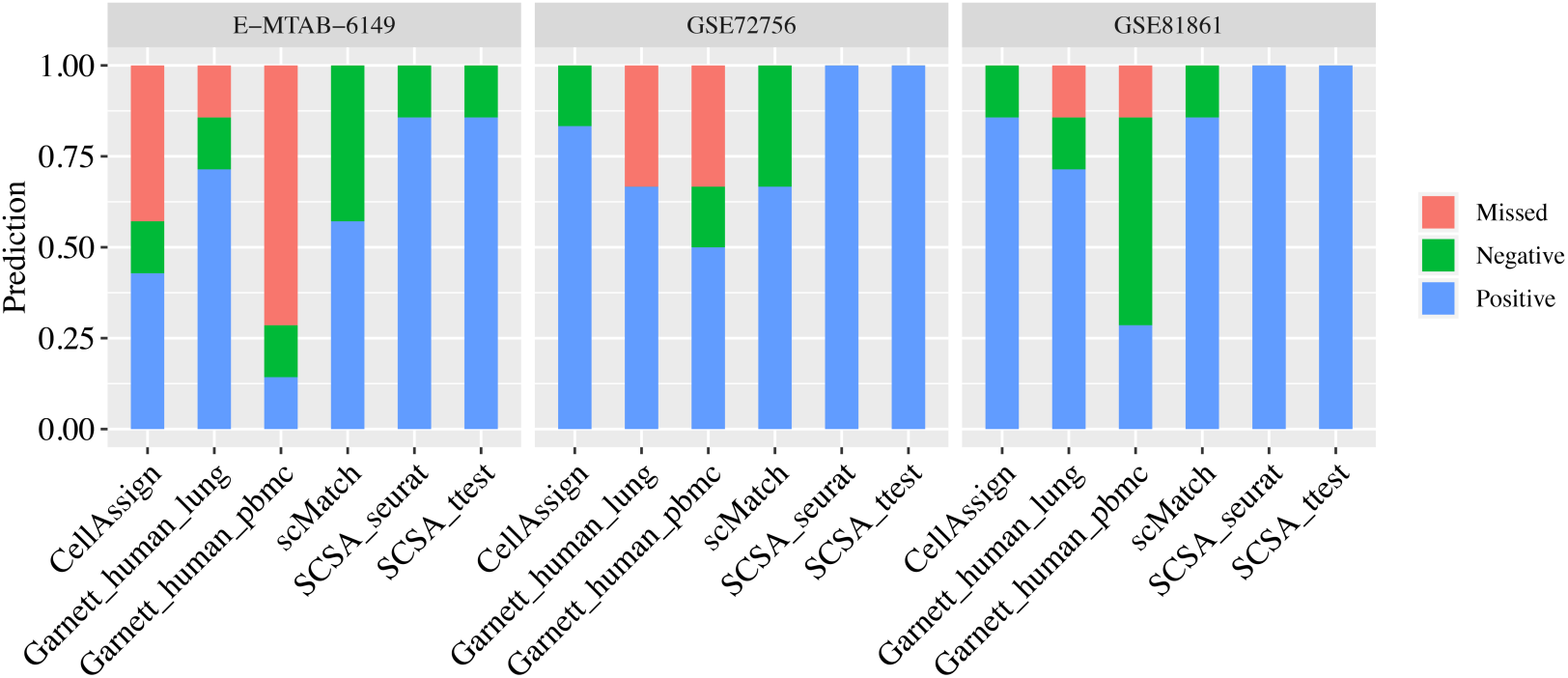
Performance of SCSA in comparison with other methods (scMatch, CellAssign and Garnett) based on 3 known cell type datasets. Dataset identity was labeled on top of the panel with cluster numbers in brackets. For legends, “Positive” meant percentage of correctly predicted clusters, while “Negative” meant incorrectly predicted clusters and “Missed” meant predictions with uncertain cell types.

**Figure 3.**
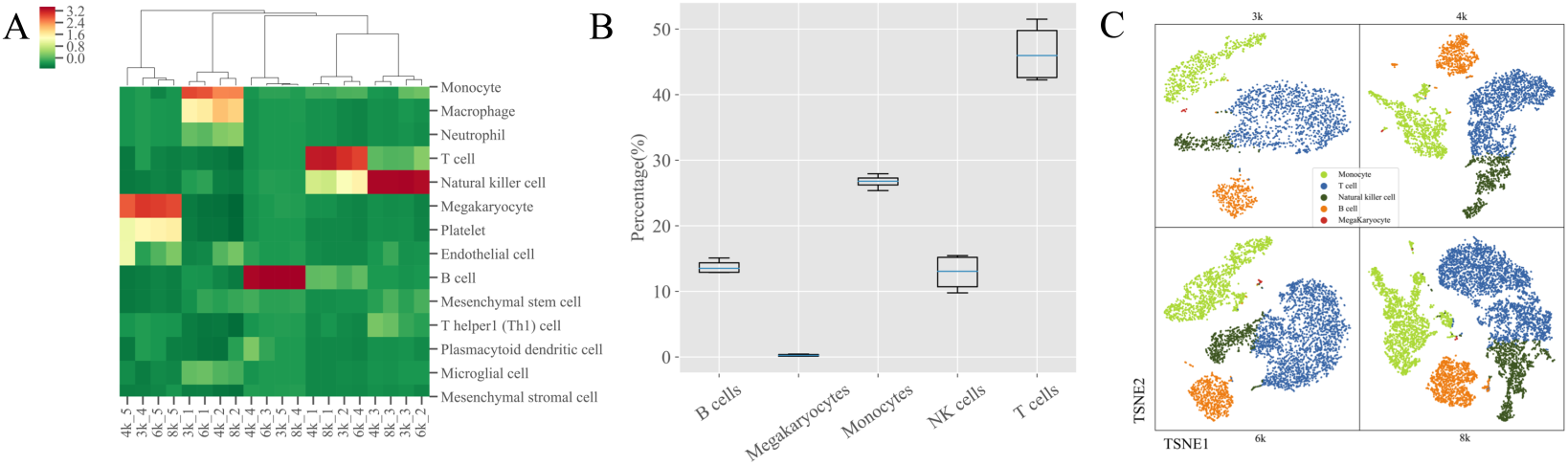
Cell components of PBMCs predicted by SCSA. A. Clustering of uniform scores of the top 5 predicted cell types in 4 PBMCs datasets by SCSA. Each column stands for one cluster of 4 PBMCs datasets and each row stands for one cell type. Uniform scores were normalized using the z-score method to make clusters comparable. B. Percentages of 4 different cell types in 4 PBMCs datasets based on SCSA’s prediction. C. Five cell types plotted by t-SNE based on the prediction of SCSA for 4 PBMCs datasets.

To further demonstrate the robustness of SCSA over the five annotated cell types (monocytes cells, T cells, NK cells, megakaryocytes cells, and B cells), we compared their abundance in each cluster using the four PBMCs datasets. As shown in Figure 3B, the percentages of cell numbers for five cell types annotated by SCSA remained stable across these datasets. T cells occupied half of the PBMCs, and monocytes cell represented another 25%, B cells and NK cells had similar levels, while megakaryocytes cell has the lowest number among all the five cell types. Meanwhile, as shown in Figure 3C, SCSA can predicted the five cell types consistent with the reference information of 4 PBMCs.

In addition, to test the GO module in SCSA, which is designed to give some clues on cluster cell function in addition to its label, we collected gene expression profiles of primary cells and cell lines from FANTOM5+SingleR dataset, which contains both human and mouse cell types (Table S8). For cell types with 10 or more samples, SCSA achieved a 60% (15 of 25) and a 57% (12 of 21) accuracy for human and mouse data respectively. For cell types containing more than 40 samples, the accuracy of SCSA were improved to 73% (8 of 11) for human data and achieved 56% (5 of 9) for mouse data, respectively (Table S2). For the “Aortic smooth muscle cell” cell type, which was not identified correctly due to lack of evidence in the CellMarker reference database, a GO analysis step in SCSA revealed a functional enrichment in the term “extracellular matrix structural constituent”, suggesting its role in regulating cell shape and cross-talk with extracellular matrix (Owens et al., 2004).

### Comparison to other cell type annotation tools

We compared the performance of SCSA with three other tools (scMatch (Hou et al., 2019), CellAssign (Zhang et al., 2019a) and Garnett (Pliner et al., 2019a)) using 3 known cell type datasets. Since scMatch, CellAssign and Garnett annotate each of the single cells instead of cell clusters, all cells were first annotated and a cell type with the maximum occurrence in the cluster was defined as the final predicted result of the cluster (Table S2-S8). As showed in Figure 2, for the three human tumor datasets (GSE72756, GSE81861 and E-MTAB-6149), SCSA achieved the highest accuracy. scMatch owned the similar accuracy with Garnett when using the predefined lung classifier (Garnett_human_lung) on the three human tumor datasets. CellAssign had a better prediction results than Garnett on GSE72756 and GSE81861 datasets, with 83% and 86% accuracy, respectively. However, for E-MTAB-6149 dataset with more than 45,000 cells, CellAssign only yielded 43% accuracy which was lower than Garnett using pre-trained classifier from human lung (Garnett_human_lung) (71%). A possible explanation for these might be that CellAssign was not suitable to annotate cell types for large datasets. Another possible explanation for these was that E-MTAB-6149 dataset is the training-set of pre-trained classifier of Garnett.

Cell-based annotation approaches could assign multiple cell type labels to one cluster due to cell heterogeneity in the clusters. Evaluation results of CellAssign in E-MTAB-6149 illustrated that only 8 (0.1%) of B cells were correctly assigned for true B cell cluster, whereas 2,861 (51.1%) cells were mistakenly assigned as myofibroblast cells and 1,755 (31.3%) cells were not able to assign a clear cell type. In general,most predicted cell types in the cluster (>50%) from these tools were not consistent with true cell type. For example, the T cells of E-MTAB-6149 dataset was not annotated by CellAssign, the fibroblasts of GSE81861 dataset was missed by scMatch, and the natural killer cells of GSE72056 dataset were not identified by CellAssign and scMatch (Table S2, S4-S6). Moreover, Garnett failed to assign epithelial cells and T cells of E-MTAB-6149 dataset although using classifier trained from the same dataset.

## Discussion

Currently, for scRNA-seq data, cell type annotation of cell clusters after unsupervised clustering is mainly conducted manually. The limitation of the manual procedure makes it impossible to generate high-quality, reproducible, and standardized annotation results for the growing number of scRNA-seq datasets.

To address this challenge, we presented a novel tool, SCSA, for automatic annotating the cell types from single-cell RNA sequencing data, which can be applied directly on the output generated from CellRanger or Seurat. By introducing the newly developed annotation model merging DEGs and cell markers reference information to replace the manual steps, and based on the extensive evaluation of performance on 7 real known datasets, as well as comparisons with alternative methods (CellAssign (Zhang et al., 2019a), Garnett (Pliner et al., 2019a) and scMatch (Hou et al., 2019)), SCSA can perform the annotation task at a high accuracy and efficient level and have a preferentially choice on annotating cell types in tumor and rare cell type datasets.

In cell type annotation, it is usually hard to find high-quality marker genes to describe a cell cluster. A strategy is to use genes specifically expressed in a cell cluster to mark the cell type. However, using a few marker genes is often not sufficient to distinguish a cell cluster from the others. In addition, using the whole expressed gene sets may decrease the power to find the true patterns within each cell cluster. Therefore, we used DEGs in the marker gene identification step in SCSA. This step avoids the influence of ubiquitously expressed genes and collects the appropriate genes for calculating the optimal score in the annotation model. There still exist some limitations, which may influence the accuracy of cell type annotation using SCSA. First, the quantity of marker genes in these cell marker databases greatly impacted the results of cell type annotation. Since cell marker collection is far away from completion, it is possible that some cell types are unclassifiable due to the lack of appropriate markers. Specifically, this phenomenon is quite common for unknown tissues and novel sub-clusters of cells at different states. User-defined marker combinations need to be developed to solve this problem. SCSA can accept them as additional information to improve the annotation results. Second, for complex tissues such as cancer tissues, the accuracy of cell annotation is heavily relied on the clustering algorithms. Different unsupervised clustering method could have different results, especially when the cluster size is unevenly distributed in the population (Kiselev et al., 2019). In that situation, algorithms using supervised clustering may be more appropriate for cell type classification (Pliner et al., 2019b).

Compared with the results of SCSA over different datasets, SCSA exhibited a reasonable accuracy and robustness in cell type annotation. Further efforts could be made to improve the annotation ability of SCSA by taking into account more information (e.g., the more accurate information of cell marker genes, the comprehensive clustering algorithm). We believe that SCSA is an important addition to the toolbox used for single-cell studies and will greatly improve our efficiency and capacity to explore the functional potential of novel cell types.

## Author Contributions

GP proposed the method and conceived of the study. YC optimized algorithm and designed the program; YC and GP analyzed the data. YC, GP, and XW wrote the manuscript. All authors discussed the results and commented on the manuscript. All authors read and approved the final manuscript.

## Acknowledgment

We thank Dr. Fangqing Zhao for the insightful suggestions for finishing the manuscript. This manuscript has been released as a pre-print at https://doi.org/10.1101/2019.12.22.886481 (Cao et al., 2019).

